# Video gaming, but not reliance on GPS, is associated with spatial navigation performance

**DOI:** 10.1101/2023.08.10.552365

**Authors:** Emre Yavuz, Chuanxiuyue He, Sarah Goodroe, Chris Ganstrom, Antoine Coutrot, Michael Hornberger, Mary Hegarty, Hugo J. Spiers

**Affiliations:** Institute of Behavioural Neuroscience, Department of Experimental Psychology, Division of Psychology and Language Sciences, University College London, London, UK; Department of Psychological and Brain Sciences, University of California Santa Barbara, Santa Barbara, California, USA; Department of Psychology, University of Pennsylvania, Pennsylvania, USA; LIRIS - CNRS – University of Lyon, Lyon, France; Norwich Medical School, University of East Anglia, Norwich, UK

## Abstract

Recent evidence suggests that greater reliance on GPS-assisted devices is associated with poorer navigation ability. Contrastingly, studies have shown that video gaming can enhance navigation ability. While gender differences in navigation ability in favour of men are well-reported, it remains unclear if the effects of reliance on GPS and video gaming on navigation performance are influenced by gender. We investigated whether gender would influence the effect of gaming experience and reliance on GPS on navigation ability using the mobile app Sea Hero Quest, which has been shown to predict real-world wayfinding performance. Alongside navigation performance assessment we asked a series of self-report questions relating to reliance on GPS, navigation strategies and gaming experience with a group of US-based participants (n = 822, 280 men, 542 women, mean age = 26.3 years, range = 18-52 years). A multivariate linear regression model found no significant association between reliance on GPS and navigation performance for either gender. There was a significant association between weekly hours of video gaming and navigation performance which was not moderated by gender. After accounting for video game experience, gender was no longer significantly associated with navigation performance. These findings have implications for which daily activities may enhance or disrupt specific cognitive abilities. Future studies applying an interventional design and real-world navigation testing would be useful to determine whether video games playing increases navigation skill, or whether those who are good at navigating tend to play more video games.

## Introduction

Being able to maintain a sense of direction and location in order to find our way in different environments is a fundamental cognitive function that relies on multiple cognitive domains (Newcombe et al., 2023; Spiers et al., 2023). Human navigation involves a range of processes such as planning routes, reading maps, identifying landmarks and maintaining a sense of direction (Newcombe et al., 2023; Ekstrom et al., 2018). Navigation can be based on both external representations such as maps or diagrams and internal representations derived from sensory experience (Wolbers & Hegarty, 2010). Whilst some people can easily find their way in novel environments, others are prone to getting lost (Coutrot et al., 2018; Burles & Iaria, 2020; Ekstrom et al., 2018; Wolbers & Hegarty, 2010; Weisberg & Newcombe, 2016; Weisberg et al., 2014; Spiers et al., 2023; Whittington et al., 2022). Understanding these individual differences in navigation ability is crucial given that deficits in navigation may constitute the earliest signs of Alzheimer’s disease (Coughlan et al., 2018), and are apparent in other conditions such as Traumatic Brain Injury (Seton et al., 2023). Understanding these individual differences in navigation will also advance the field of spatial cognition at large (Newcombe et al., 2023; Spiers et al., 2023).

Being able to create a valid test of navigation that accounts for the wide variation in navigation performance is challenging, given the large sample sizes needed and the high levels of environmental manipulation and experimental control required in standard research settings (Newcombe et al., 2023). With the recent evolution of widespread touch-screen technology on both tablet and mobile devices and virtual reality (VR), our team developed a series of navigation tests in the form of a mobile video game app Sea Hero Quest (SHQ)(Spiers et al., 2023). SHQ has since been used to test the navigation ability of 4 million people globally, has good test-retest reliability, and is predictive of real-world navigation performance (Coutrot et al., 2018; Coutrot et al., 2019; Coughlan et al., 2020; Spiers et al., 2023). Studies using SHQ in healthy participants so far have shown that the often-reported superior navigation performance in men may be partly accounted for by gender inequality (Coutrot et al., 2018), that participants are better at navigating in environments that are similar in topology to the environment they grew up in (Coutrot et al., 2022a) and that spatial navigation performance is greatest in those who sleep for 7 hours a night (Coutrot et al., 2022b). On a clinical level, studies so far have shown that SHQ performance can be used to classify spatial impairments in healthy participants with a high risk of Alzheimer’s Disease (Coughlan et al., 2019), to detect AD patients most prone to disorientation (Puthusseryppady et al., 2022), and to detect spatial navigation deficits in those with traumatic brain injury (Seton et al., 2023).

With the rise in technological devices, reliance on GPS-based systems to aid human navigation has become significantly more prevalent (He & Hegarty, 2020; McKinlay, 2016). Numerous studies have shown that using GPS-based systems may be detrimental to human navigation performance, as assessed using self-report questionnaires (He & Hegarty, 2020; Ishikawa, 2018) computerised and real-world studies (Dahmani & Bohbot, 2020; Fenech et al., 2010; Gardony et al., 2018; Hejtmanek et al., 2018; Ishikawa et al., 2008; Kippel et al., 2010; Parush et al., 2007; Ruginski et al., 2019; Schelton et al., 2013; Schwering et al., 2017; Willis et al., 2008). For example, the negative effect of spatial anxiety on one’s self-reported sense of direction was mediated by a greater reliance on GPS (He & Hegarty, 2020), and that greater reliance on GPS was significantly associated with poorer spatial memory when participants were required to find their way in an environment they had learnt without using GPS (Dahmani & Bohbot, 2020).

While reliance on GPS has been found to be associated with poorer navigation performance, video game playing has been associated with better cognitive performance in a number of studies (Anguera et al., 2013; Brilliant et al., 2019; Gleich et al., 2017; Green & Bavelier., 2003; Haier et al., 2009; Hong et al., 2023; Hill et al., 2017; Kuhn et al, 2013; Lee et al., 2012; Leitao et al., 2020; Lorenz et al., 2015; Martinez et al., 2012; Rosser et al., 2007). Video game-based interventions on VR and mobile app-based platforms have also shown promise in detecting those with neurodegenerative disorders such as Multiple Sclerosis (van der Ham et al., 2022) and Alzheimer’s Disease (Poos et al., 2021; Innamuri et al., 2021), as well as for brain rehabilitation training in those with cognitive decline (Brugada-Ramentol et al., 2022). In healthy participants, both performance on memory-based tasks associated with-and neural plasticity within the hippocampus, a key region of the navigation network, has been shown to increase as a consequence of playing both VR-and mobile app-based video games (Clemenson et al., 2019; Kuhn et al., 2013; Krokos et al., 2018; Martin et al., 2022; Stark et al., 2021; West et al., 2018). Additionally, playing video games has been shown to lead to improvements in other cognitive domains aside memory that are related to spatial navigation, including spatial attention, mental rotation, spatial perspective taking and cognitive mapping (Feng et al., 2007; Green & Bavelier, 2003; Lin et al., 2020; Mclaren-Gradinaru et al., 2020; Murias et al., 2016).

Numerous studies using SHQ have shown that men have superior navigation ability to women (Coughlan et al., 2019, 2020; Coutrot et al., 2018, 2019, 2022a, 2022b; Puthusseryppady et al., 2022; Seton et al., 2023; Spiers et al., 2023; West et al., 2023), corroborating similar findings in the wider literature (Narazeth et al., 2019). However, other studies have shown that sex differences in spatial cognition are no longer apparent when accounting for prior video game experience, where men lose their well-reported advantage in mental rotation ability (Feng et al., 2007; Liu et al., 2020; McLaren-Gradinaru et al., 2023). Men tend to spend more hours playing video games than women (Dindar, 2018; Entertainment Software Association, 2010, 2022; Leonhardt & Overa, 2021; Lucas & Sherry, 2004; McLaren-Gradinaru et al., 2023; Ogletree & Drake, 2007; Quaiser-Pohl et al., 2006). It therefore remains a possibility that the superior navigation ability in men is a result of their greater video gaming experience.

The navigation strategy one uses when playing video games may also play a role in influencing the effect of gaming on spatial navigation performance. Accordingly, it was shown that the effect of playing video games was found to be detrimental or beneficial depending on the navigation strategy the player used, as well as the genre (West et al., 2018). In a reciprocal fashion, video gaming experience can also influence the navigation strategy used during spatial navigation tasks, such that those who played video games reported using more efficient navigation strategies for orientation such as using cognitive maps or learned routes (Murias et al., 2016). Additionally, men are more likely to play action, simulation and role-playing-based video games that have a strong navigational component (Dindar et al., 2018; Lucas & Sherry, 2004; Quaiser-Pohl et al., 2006), suggesting that their greater video game experience may account for their greater use of more efficient navigation strategies, as reported in the literature (Boone et al., 2018). A key questionnaire that has been used to assess everyday use of navigation strategies is the Navigation Strategies Questionnaire (NSQ)(Brunec et al., 2019). Studies using this questionnaire have shown that the navigation strategy one uses influences navigation performance (Brunec et al., 2019), whilst our recent study using SHQ and the NSQ have found navigation strategy is not significantly associated with navigation performance (West et al., 2023). Thus, the extent to which one’s choice of navigation strategy may account for navigation performance remains unclear.

Bringing together these findings, we hypothesised that those with a greater reliance on GPS would show worse navigation performance, whilst those with more hours of video gaming experience would show better navigation performance on SHQ. Secondly, we hypothesised that the association between video game experience and navigation performance would be stronger in males. Thirdly, we hypothesised that gender differences in navigation performance would be mitigated after accounting for video game experience. As the extent of gender differences in the reliance on GPS remains unclear from the literature (for preliminary results, see Miola et al., 2023), we did not make a specific hypothesis about how the effect of reliance on GPS on navigation performance would be influenced by gender. Given the broader impact of technology use on cognitive function (Small et al. 2020; van der Schuur et al., 2015), we explored whether there would be a significant interaction between video game experience and reliance on GPS on navigation performance. Finally, we explored whether the NSQ would be a reliable measure of the navigation strategies one uses in everyday life using psychometric analysis, in order to control for any differences in the navigation strategies participants used when assessing their navigation performance.

## Methods

### Participants

903 participants living in the US aged 18 and above (309 men, 575 women, other, mean age = 27.0 years, SD = 8.0 years, range = 18-66 years, mean number of years of formal education = 16.1 years) were recruited using the Prolific database (www.prolific.co, 2023) and reimbursed for their time. Ethical approval was obtained from the University College London Review Board conforming to Helsinki’s Declaration. All participants provided informed written consent. We removed 19 participants who selected ‘other’ for gender as we believed it would be hard to make any informative interpretation of any findings with this limited sample size. We then further identified outliers in this remaining sample of participants using Mahalanobis’ Distance, a method shown to have high sensitivity and specificity and a minimal change in bias in simulated and real datasets when removing outliers based on questionnaire data compared to other methods (Curran, 2016; Zijlstra et al., 2011). 79 participants were removed as outliers. This resulted in a final sample size of 822 participants (367 men, 455 women, mean age=26.3 years, SD=6.7 years, range=18-52 years, mean number of years of formal education=16.1 years). Demographic information for the final sample is summarised in Table 1. Data analysis was completed using R studio (version 1.4.1564) and Python (version 3.9.12).

**Table 1.** Summary of basic descriptive statistics for the variables included in the model. These were: weighted wayfinding distance, age, weekly hours of video game use on all devices, weekly hours of phone use, for men and women separately, and across gender. Mean and standard deviation (SD) values are shown. For weighted wayfinding distance, the raw (not z-scored) wayfinding distance is shown. VR-m = virtual reality metres.

|  | Women | Men |
| --- | --- | --- |
| n (%) | 455 (55.4%) | 367 (44.6%) |
| Weighted wayfinding distance (VR-m) | 11.76 (3.14) | 10.69 (3.69) |
| Age in yrs | 24.80 (5.88) | 28.05 (7.20) |
| Average GPS reliance score (1-5, Max. 5) | 3.72 (0.69) | 3.41 (0.78) |
| Weekly hours of phone use | 36.07 (23.06) | 28.90 (21.84) |
| Weekly hours of video game use on all devices | 4.78 (5.36) | 10.13 (6.33) |

### Statistical Power Analysis

A power analysis was conducted using G* power (Erdfelder et al., 1996). 822 participants were sufficient to achieve a small effect size (Cohen’s f2 = 0.03), at an alpha threshold of 0.05 with 95% power (Selya et al., 2012).

### Experimental Procedure

#### Self-report questionnaires

To characterise the use of navigation strategies, the Navigation Strategies Questionnaire (NSQ) was used (Brunec et al., 2019). This questionnaire has fourteen items and assesses the degree to which an individual relies on map-based strategies when navigating in the real world. Questions refer both to the strategies one uses when navigating (e.g., “when planning a route, do you picture a map of your route or do you picture scenes of what you will see along the way?”) and one’s navigation ability (e.g., “do you find it easy to read and use maps?”).

To characterise one’s reliance on GPS, the GPS reliance scale was used (Dahmani and Bobot, 2020). This scale has seven items and assesses the frequency at which people have relied on GPS in different situations within the past month (e.g., “How often do you use GPS to travel new routes to a previously visited destination?”). The average score across questions was calculated for each participant.

To characterise video game use, participants were asked to indicate the number of hours per week spent playing video games on all devices per week, as well as specifically on a phone or tablet, the number of hours of phone use per week, their most commonly played video game genre and the video game platform they used most often. Participants were also asked to report their age, gender and highest education level. Please see Supplementary Materials for the full set of questionnaires.

#### Sea Hero Quest Task

Sea Hero Quest (SHQ) is a VR-based video game for mobile and tablet devices which requires participants to navigate through a three-dimensional rendered world in a boat to search for sea creatures in order to photograph them, with the environment consisting of ocean, rivers and lakes (Coutrot et al., 2018; Spiers et al., 2023). Although navigational abilities on SHQ have been assessed using Wayfinding, Path Integration and Radial Arm Maze measures, we focused on wayfinding in this study (Figure 1). We asked participants to play 5 levels -levels 1, 11, 32, 42 and 68 - where level 1 was a tutorial level designed to assess one’s ability to control the boat, whilst the latter 4 levels were wayfinding levels. We selected these specific levels as they showed the greatest effect sizes when investigating the effect of one’s home environment on wayfinding performance in a previous study (Coutrot et al., 2022a). The wayfinding levels increased in difficulty from level 11 to 68, with the difficulty of a given level based on the number of goals and how far apart they were from each other (see Yesiltepe et al., 2023). Participants were required to play all 5 levels, where playing a given level would unlock the next level. For each participant, we quantified the wayfinding distance, defined as the euclidean distance travelled between each sampled location in pixels, for levels 11, 32, 42 and 68 separately. We then divided the wayfinding distance in each level by the wayfinding distance in level 1 to control for the effects of gaming experience on navigation performance. To control for the difference in wayfinding distance between levels, we then z-scored the distances within each level and averaged these across the levels. This resulted in each participant having a z-score which represented their wayfinding distance across the 4 levels. This z-score was referred to as the weighted wayfinding distance. A shorter weighted wayfinding distance indicated better navigation performance (i.e., a more efficient route to the goal).

**Figure 1.**
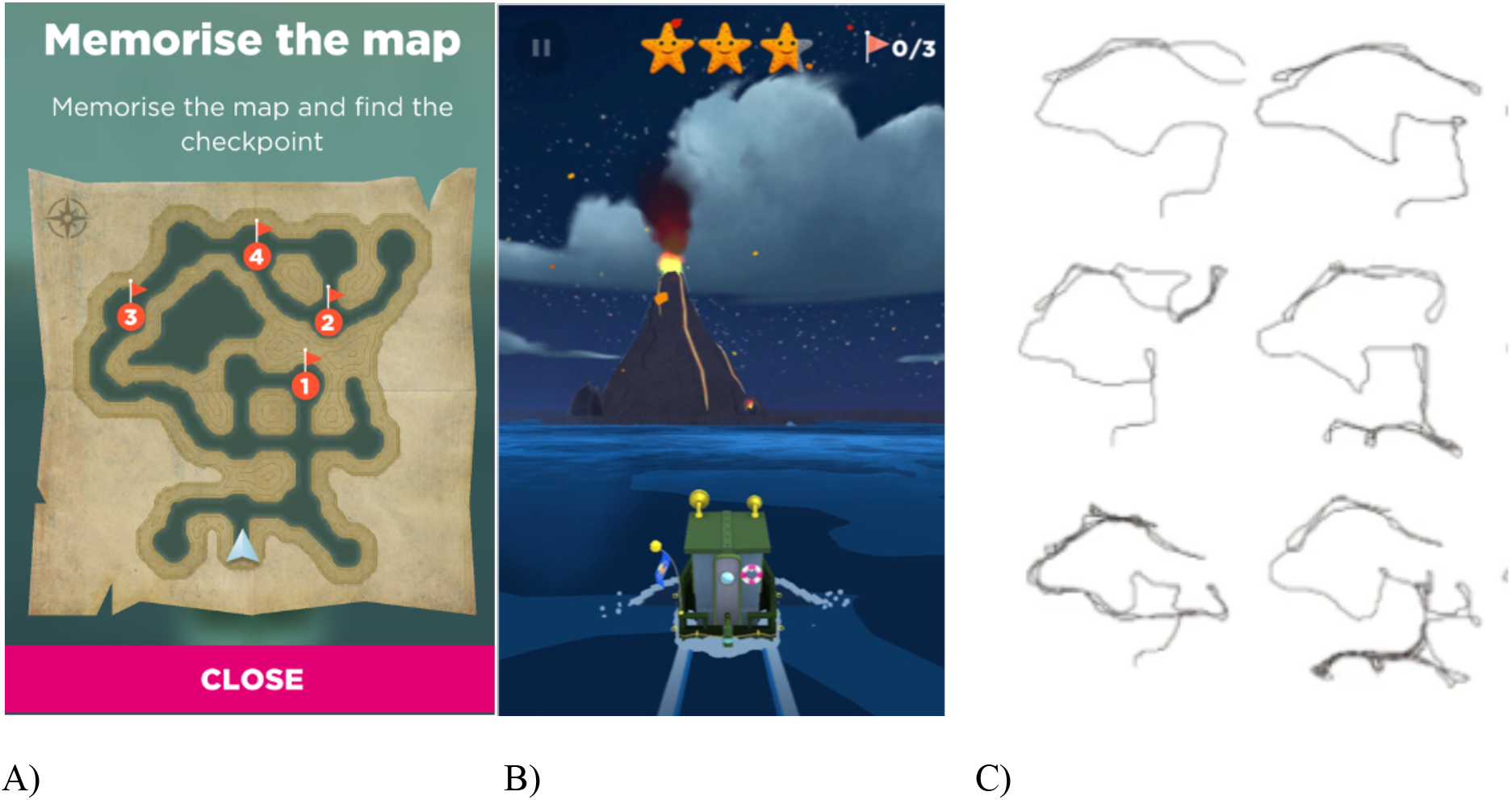
Outline of the wayfinding task. (A) At the start of each level presented participants were presented with a map indicating the goals which they had to navigate to in the order indicated, where ‘1’ indicates goal number 1 etc. (the map from level 42 is shown as an example). Level 1 (not shown) provided one goal and a simple river to reach the goal as training. (B) Participants selected to close the map by pressing ‘close’, at which point the participant had to start navigating to the goals. First-person view of the environment is shown where the participant tapped left or right of the boat to steer it. (C) Examples of individual player trajectories (level 42) from the start location to the final goal. Trajectories are ordered by performance, with the top left providing the best performance (shortest trajectory length), through to bottom right who has the worst performance (longest trajectory length). Adapted from Coutrot et al. (2018).

## Data Analysis

As the NSQ includes questions relating to a participants’ navigation ability and strategy, we conducted an exploratory factor analysis to determine whether the NSQ could be explained by a strategy and an ability subscale. McDonald’s omega factor saturation, split-half permutation testing and Chronbach’s alpha reliability testing was performed to assess the internal reliability of the NSQ (Parsons et al., 2019). For the split-half permutation reliability analysis we used an odd-even split with 5000 permutations, where a higher Spearman-Brown correlation value indicates a higher internal reliability. A higher value for Chronbach’s alpha indicates a higher internal reliability of a given questionnaire (Tavakol & Dennick, 2011). For McDonald’s omega factor saturation testing, a higher value for total omega indicates a higher internal reliability of a given questionnaire (Hayes & Coutts, 2020). We also performed these same tests on the GPS reliance scale and the weighted wayfinding distances across the wayfinding levels (Supplementary Table S8).

To measure the associations between self-report questionnaire responses and wayfinding distance, we first conducted Pearson’s correlations between age, average GPS reliance score, hours of video game use per week on all devices and weighted wayfinding distance to explore the strength of the associations between each of these variables. We conducted point-biserial correlations between each of these variables and gender given that gender was a binary variable.

We then conducted a multivariate linear regression model to analyse the effects of gender, average GPS reliance score and hours of video game use per week on all devices on wayfinding distance. We removed the fifth question of the GPS reliance scale - “You usually travel a specific route to go to your friend’s house. This time, you think you may get there faster by taking a different route. How often do you take new routes to travel to places you have visited before?” - as we did not believe this was assessing reliance on GPS. To understand how the effect of gender on navigation performance was influenced by GPS reliance and video game experience, we included a GPS reliance x gender and an hours per week video game use x gender interaction terms in the model. We also included a GPS reliance x hours of video game use per week interaction term to determine whether the effects of reliance on GPS would interact with the effects of video gaming on navigation performance. Age and highest level of education were included in the model as covariates given their previous association with wayfinding distance (Coutrot et al., 2022a). Given that SHQ is a mobile app-based video game, we also included hours of weekly phone use as a covariate. We then conducted a second variation of this model without including weekly hours of video gaming as a predictor variable, in order to determine whether gender was significantly associated with navigation performance when video gaming experience was not accounted for. We also conducted a third variation of this model to explore whether there was an effect of video games playing on the weighted score from the tutorial level (level 1) which does not test navigation, but might be mastered more efficiently by those familiar with video games. Finally, as a supplementary analysis, we conducted a fourth variation of this model where the most commonly played video game genre and the most commonly used video game platform were included as variables of interest. For this model, we also excluded the 101 participants who did not play video games. The purpose of this model was to explore whether the associations found to be significant in the main model would remain significant when accounting for the video game genre most commonly played and the video game platform most commonly used (see Supplementary Materials).

As a check for multicollinearity, we calculated the adjusted generalised variance inflation factor (*GVIF*) value for each predictor variable included in the model, which is equal to the Variance Inflation Factor (*VIF*) values for each predictor variable divided by the number of categories for each categorical predictor variable:*GVIF* = *VIF* ^[^*^1^*^/(*2*∗&’)]^. The *GVIF* value was scaled to account for differences in the number of degrees of freedom for each of the predictor variables in the model, producing an adjusted *GVIF* value: *adjusted GVIF* = *GVIF ^[^*^1^*^/(*2*∗&’)]^*. For cases where a model contained a categorical variable with more than two categories, the *GIVF* ^[^*^1^*^/(*2*∗&’)]^ value is displayed for each predictor variable in the model. In cases where the model contains no predictor variable with more than two categories, the *VIF* value is displayed. A *VIF* or *GVIF* ^[^*^1^*^/(*2*∗&’)]^ value of <5 indicates that multicollinearity is not a concern (Fox and Monet, 1992; Kim, 2019). Post-hoc t-tests bonferroni-corrected for multiple comparisons to control for type 1 errors were carried out where main effects and interactions were significant.

## Results

### Gender differences in video gaming, reliance on GPS, age and phone use

Unpaired t-tests indicated that men had significantly greater weekly hours of video game use on all devices (*t =* -12.89*, p* < 0.001) and had a significantly lower average GPS reliance score (*t =* 5.85*, p <* 0.001) than women. Men were also significantly older (*t =* -6.98*, p <* 0.001, 28.1 yrs vs 24.8 yrs), had significantly lower weekly hours of phone use (*t =* 4.57*, p* < 0.001) and had a significantly shorter wayfinding distance than women (*t =* 4.43*, p <* 0.001), indicating more efficient wayfinding. All these results were corrected for multiple comparisons using Bonferroni’s method (Table 1).

### Bivariate analysis - reliance on GPS was not significantly associated with wayfinding distance, whilst hours of weekly video gaming was

Pearson’s correlations indicated that the average GPS reliance score was not significantly associated with wayfinding distance (*r* = 0.01, *p* = 0.794, *CI* = [-0.06, 0.08]). Hours of video game use per week on all devices was significantly associated with wayfinding distance (*r* = -0.23, *p* < 0.001, *CI* = [-0.29, -0.16]), when controlling for multiple comparisons (p < 0.008, bonferroni-corrected with 6 comparisons)(Figure 3). See Figure 3 for the full set of correlations between all variables.

**Figure 3.**
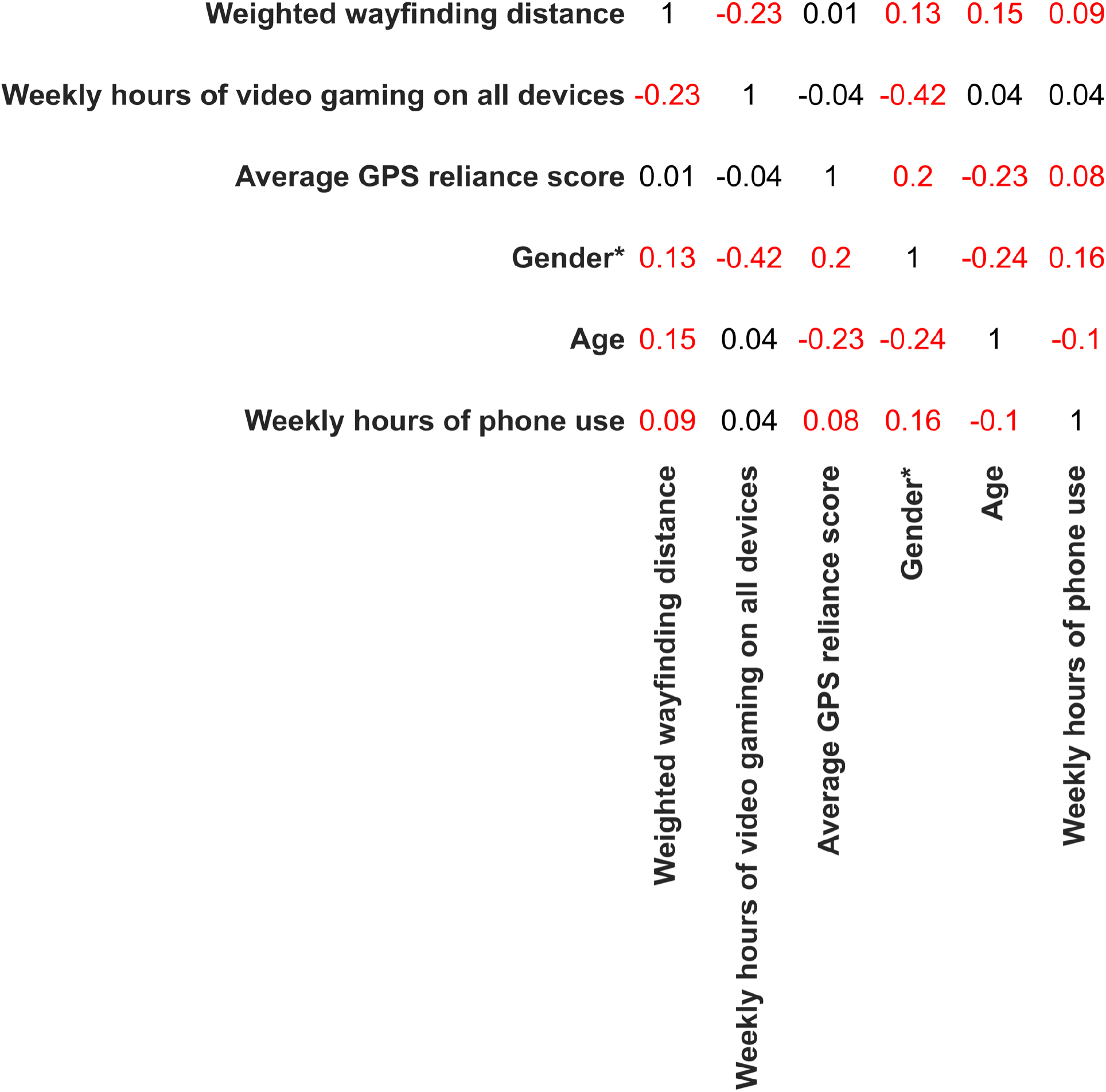
Pearson’s correlation coefficients for the associations between each of the variables included in the model. *Point-biserial correlation coefficients were calculated for the associations between gender and each of the other variables given that gender was a binary variable (man/woman). Values highlighted in red indicate significant associations.

### Multivariate analysis - reliance on GPS was not significantly associated with wayfinding distance, whilst hours of weekly video gaming was, when controlling for other variables

After examining these initial correlations, we then conducted a linear multivariate regression model to verify these associations when controlling for the associations of other variables with wayfinding distance. In this analysis, GPS reliance score, hours of video game use per week were the key variables of interest, whilst age, gender and hours of phone use per week were covariates. Interactions between gender and both average GPS reliance score and hours of video game use per week on all devices were included in the model (Table 3). Before running the regression, as a check for multicollinearity, we calculated the *VIF* value for each predictor variable in the model. This revealed that all the predictor variables had a *VIF* value that was less than 5, indicating that multicollinearity was not a concern (Supplementary Table S1).

**Table 3.**
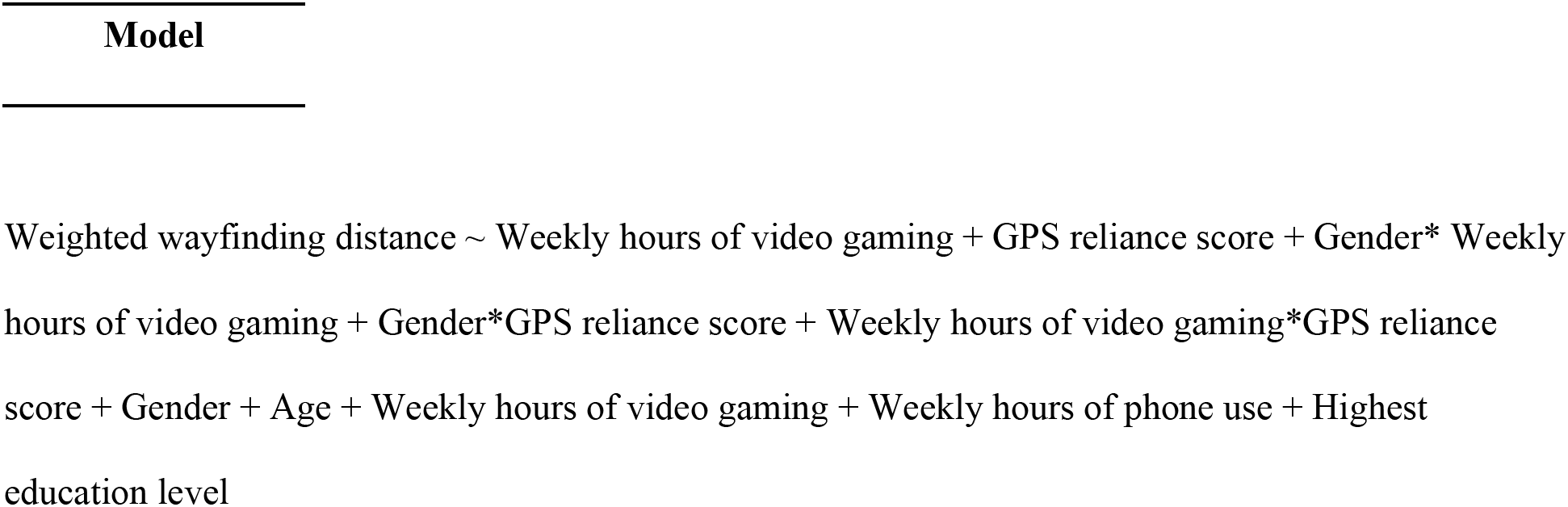
Model specification.

| Model |
| --- |
| Weighted wayfinding distance ~ Weekly hours of video gaming + GPS reliance score + Gender* Weekly hours of video gaming + Gender*GPS reliance score + Weekly hours of video gaming*GPS reliance score + Gender + Age + Weekly hours of video gaming + Weekly hours of phone use + Highest education level |

The outputs from the multivariate linear regression model are as follows:

### Main variables of interest

#### Reliance on GPS

Contrary to our hypotheses, average GPS reliance score was not significantly associated with wayfinding distance (*β* = -0.03, *f2* = < 0.001, *p* = 0.350, *CI* = [-0.08, 0.03])(Table 4 and Figure 4). Moreover the interaction between average GPS reliance score and gender did not reach significance (*β* = 0.07, *f2* = <0.001, *p =* 0.08*, CI =* [-0.01, 0.15])(Table 4 and Figure 4 indicating that the non-effect of GPS held for both men and women.

**Figure 4.**
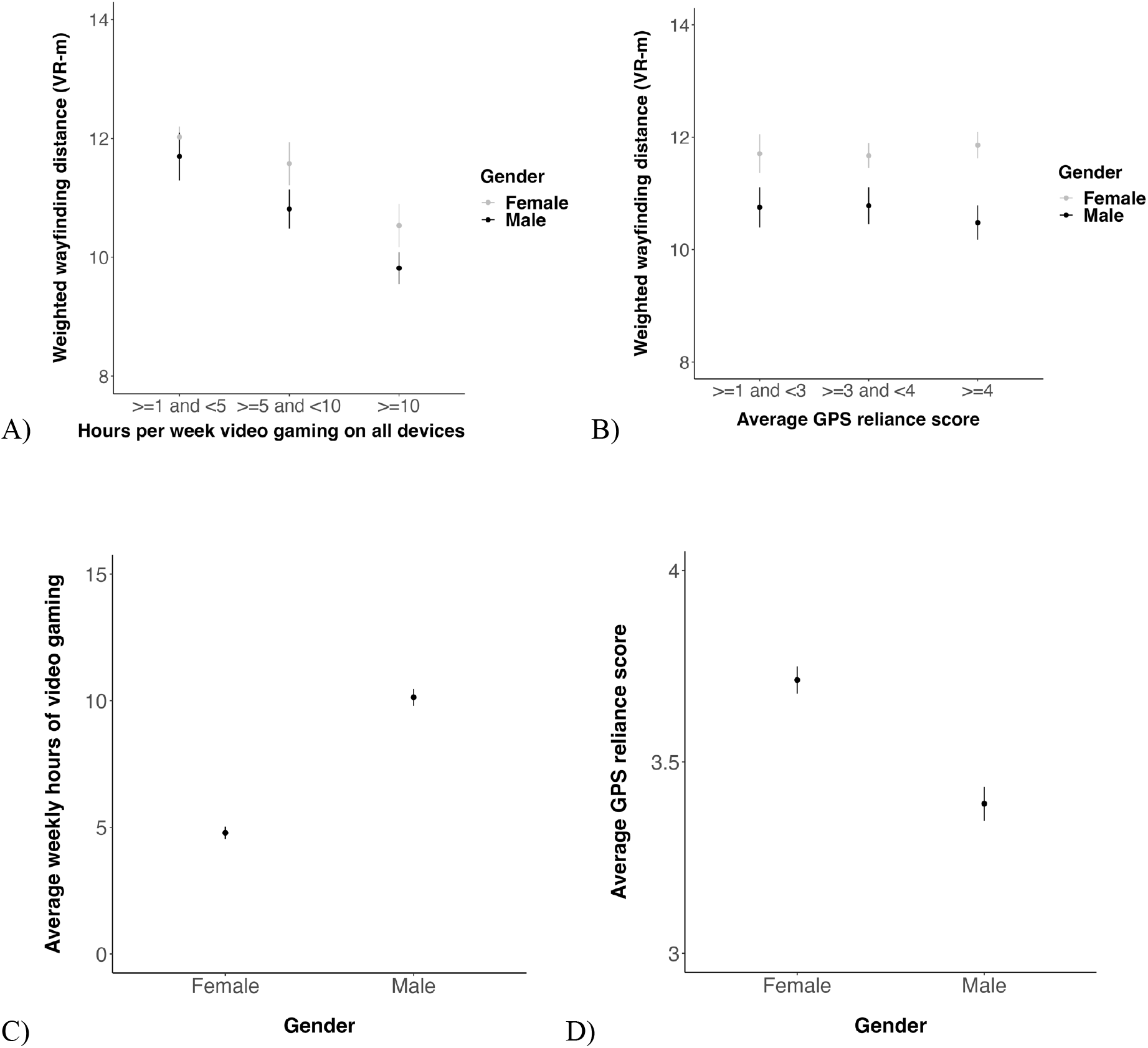
Associations between both weekly hours of video gaming and average GPS reliance scale scores and weighted wayfinding distance across gender and in men and women separately. (A-D) Data points represent the mean wayfinding distance across game levels across participants. Bars represent one standard error above and below the mean wayfinding distance across game levels across participants. VR-m = virtual reality metres.

**Table 4.** Output of the model predicting weighted wayfinding distance based on GPS reliance and video game experience. P-values for the significant associations are highlighted in bold.

| Variable | $\beta$ | 95% CI | $t$ | $p$ | sig | $f^2$ |
| --- | --- | --- | --- | --- | --- | --- |
| (Intercept) | -0.17 | [-0.26, -0.09] | -4.21 |  |  |  |
| Age | 0.10 | [0.06, 0.14] | 4.68 | **<0.001** | *** | 0.02 |
| Male gender | 0.07 | [-0.02, 0.15] | 1.50 | 0.134 |  | 0.03 |
| Average GPS reliance score | -0.03 | [-0.08, 0.03] | -0.94 | 0.350 |  | <0.001 |
| Weekly hours of phone use | 0.06 | [0.02, 0.10] | 3.04 | **0.002** | ** | 0.01 |
| Weekly hours of video gaming on all devices | -0.12 | [-0.17, -0.06] | -4.03 | **<0.001** | *** | 0.04 |
| Highest education level achieved | 0.06 | [-0.02, 0.14] | 1.43 | 0.153 |  | <0.001 |
| Gender*Average GPS reliance score | 0.07 | [-0.01, 0.15] | 1.73 | 0.084 | . | <0.001 |
| Gender*Weekly hours of video gaming on all devices | -0.01 | [-0.09, 0.08] | -0.15 | 0.883 |  | <0.001 |
| Average GPS reliance score*Weekly hours of video gaming on all devices | 0.04 | [<0.001, 0.08] | 1.89 | 0.060 | . | <0.001 |

#### Hours of weekly video gaming on all devices

As predicted, hours per week video game use on all devices per week was significantly associated with wayfinding distance (*β* = -0.12, *f2* = 0.04, *p* < 0.001, *CI =* [-0.17, -0.06]) (Table 4 and Figure 4). Post-hoc t-tests revealed that those playing >=10 hours of video games per week on all devices were significantly better navigators than both those playing >=5 hours but <10 hours and those playing <5 hours per week, as indicated by the shorter wayfinding distance in the former group (*p* < 0.017, bonferroni-corrected with 3 comparisons)(Table 5). The interaction between the hours of video game use and gender did not reach significance (*β* = -0.01, *f2* = < 0.001, *p* = 0.883, *CI =* [-0.09, 0.08]), nor did the interaction between the hours of video game use on all devices per week and average GPS reliance score (*β* = 0.04, *f2* = <0.001, *p* = 0.06, *CI =* [<0.001, 0.08])(Table 4 and Figure 4).

**Table 5.**
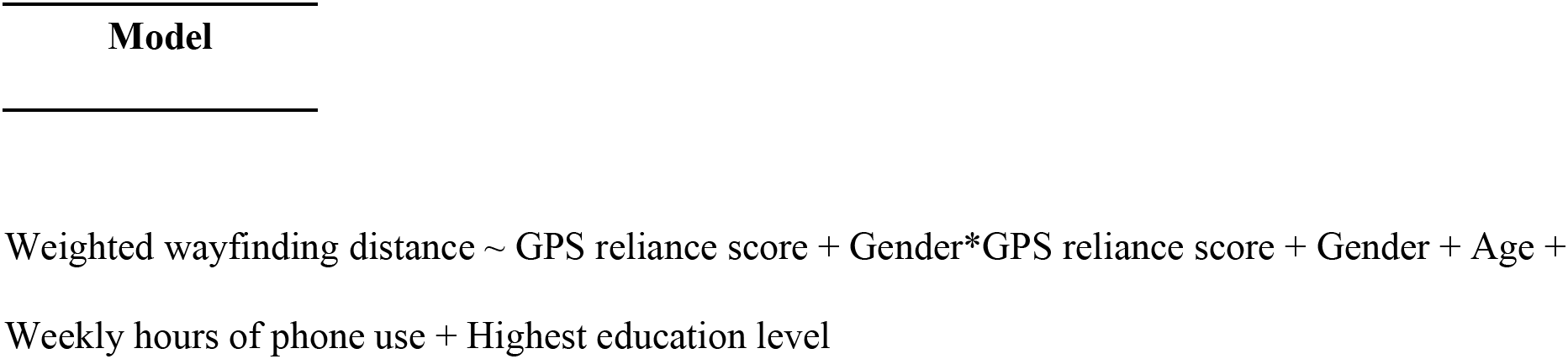
Model specification.

#### Covariates

Please see Supplementary Figures S2A-D for visualisations of the associations between each of the covariates with wayfinding distance.

#### Multivariate analysis - gender was significantly associated with navigation performance when video game experience was not accounted for

We next conducted a second linear multivariate regression model to verify whether gender would be significantly associated with navigation performance when video game experience was not accounted for. For this model, we used the same predictor variables as in the model above except that we removed video game experience as a predictor variable (Table 5). Like with the previous model, all the predictor variables had a *VIF* value that was less than 5, indicating that multicollinearity was not a concern (Supplementary Table S2).

As predicted, gender was significantly associated with wayfinding distance in this model (*β* = 0.17, *f2* = 0.03, *p* < 0.001, *CI =* [0.09, 0.25]) (Table 6).

**Table 6.** Output of the model predicting weighted wayfinding distance without including video game experience as a predictor variable. P-values for the significant associations are highlighted in bold.

| Variable | $\beta$ | 95% CI | *t* | *p* | sig | *f*<sup>2</sup> |
| --- | --- | --- | --- | --- | --- | --- |
| (Intercept) | -0.25 | [-0.33, -0.17] | -6.46 |  |  |  |
| Age | 0.1 | [0.06, 0.14] | 4.74 | **<0.001** | *** | 0.02 |
| Male gender | 0.17 | [0.09, 0.25] | 4.2 | **<0.001** | *** | 0.03 |
| Average GPS reliance score | -0.01 | [-0.06, 0.04] | -0.38 | 0.71 |  | <0.01 |
| Weekly hours of phone use | 0.04 | [0.01, 0.08] | 2.24 | **0.03** | * | 0.01 |
| Highest education level achieved | 0.09 | [0.01, 0.17] | 2.13 | **0.03** | * | 0.01 |
| Gender*Average GPS reliance score | 0.04 | [-0.04, 0.11] | 0.91 | 0.36 |  | <0.01 |

#### Complementary analysis

### Examining how tutorial level performance (Wayfinding distance on level 1 of SHQ) relates to measures of interest

When running the same multivariate model above for level 1 wayfinding distance only, with the most commonly used video game genre as a covariate, weekly hours of video gaming experience was not significantly associated with weighted wayfinding distance (*β* = -0.03, *f2* = <0.001, *p* = 0.550, *CI* = [-0.12, 0.06])(Supplementary Table S8).

#### Psychometric testing of the Navigation Strategies Questionnaire (NSQ) reveals a poor internal reliability

Exploratory factor analysis showed that the NSQ could only be poorly explained by a strategy and ability subscale. McDonald’s Omega, Cronbach’s alpha and Split-half permutation testing also demonstrated that it had poor internal reliability (Supplementary Table S9). We therefore decided not to include it in our multivariate linear model.

## Results Summary

Taken together, there are three key findings. Firstly, average GPS reliance score was not significantly associated with navigation performance, nor did it show a significant interaction with gender to influence navigation performance. Secondly, greater weekly hours of video gaming was associated with better navigation performance, although the strength of this association did not differ by gender. Thirdly, when accounting for weekly hours of video gaming, gender differences in navigation performance were no longer significant.

## Discussion

Using a virtual navigation test embedded in a video game and a set of questionnaires, we examined whether reported reliance on GPS or reported video games playing was associated with navigation performance. We found no significant association between average GPS reliance scores and wayfinding distance, contradicting our hypothesis and in contrast with previous findings (e.g., Dahmani & Bohbot, 2020; He & Hegarty, 2020). However, we did find an association between video games playing and wayfinding distance consistent with our predictions. We discuss these key results in turn.

Although our study used the same GPS reliance scale as that used by Dahmani & Bohbot, 2020 when examining reliance on GPS, and used participants within a similar age range, their navigation task differed from ours. They had participants remember the locations of objects within a radial arm maze. In our study, participants had to remember the locations of a series of checkpoints from a one-shot encoding of a cartographic map and then navigate to these checkpoints without the map available. Additionally, their study was a longitudinal study and only included 13 participants in the analysis. Thus, differences in the experimental task, study design and sample size may have accounted for differences in findings. Dahmani & Bohbot 2020 also found that a greater reliance on GPS was associated with less use of a landmark strategy during a radial arm maze task. Thus, we might have expected an association between reliance on GPS and navigation performance in SHQ because we have previously shown that landmark use in SHQ radial mazes is associated with good navigation (West et al, 2023; Garg et al., 2023). It would have been useful to probe this further with the NSQ, which seeks to measure strategy use (Brunec et al., 2019). However, given the low internal reliability of the navigation strategies questionnaire, we could not accurately assess the extent to which participants in our study may have relied on different strategies. He & Hegarty (2020) used SBSOD as a proxy for navigation, but Garg et al., (2023) showed that there was no significant association between scores on the Santa Barbara Sense of Direction Scale (SBSOD) and SHQ wayfinding performance. This may help explain why we did not find a significant association between reliance on GPS and navigation performance, whilst He & Hegarty, 2020 did.

Interestingly, not all studies have shown that reliance on GPS is detrimental to navigation performance (Brugger et al., 2019; Ercevik Somnez & Erinsel Onder, 2019; Huston & Hamburger., 2023; Kelly et al., 2022; Leshed et al., 2008; Wunderlich et al., 2022; Yan et al. 2022). For example, a recent study showed that route knowledge was not affected in those who followed turn-by-turn instructions (simulating reliance on GPS) for the duration of a route compared with those who initially followed instructions, which needed to be memorised (Kelly et al., 2022). Moreover, participants using GPS-based devices during navigation were found to form mental images of landmarks along a route to a similar extent to those who did not use such devices, and could successfully use these to navigate to future locations along the route (Ercevik Somnez & Erinsel Onder, 2019). Furthermore, another study found that dependency on GPS-based devices showed no significant association with navigation performance on a wayfinding task (Yan et al., 2022). Findings from the literature also suggest that using GPS-based devices may in fact enrich one’s ability to learn to navigate new environments, for example, a study using qualitative reports showed that using GPS during navigation can create more meaningful, enriched and genuine experiences when navigating by providing users with new tools and sources of information (Leshed et al., 2008). Studies have also found that when participants actively engage with a GPS-assisted device, the level of spatial knowledge acquired during a wayfinding task is greater than when participants rely on a fully automated GPS-assisted device (Brugger et al., 2019). Similarly performance can be improved by prompting participants to learn with the GPS rather than acting passively (Huston & Hamburger, 2023). Additionally, choosing to manually zoom or being automatically zoomed into a GPS-assisted device resulted in improved wayfinding performance in the learning and testing phases of a wayfinding task, respectively (Richter et al., 2010). Thus, the way in which participants engage with GPS-assisted devices may mediate its effect on navigation performance rather than the general reliance on GPS per se.

There was a significant association between hours of video game use and navigation performance, such that more hours of video gaming was associated with better navigation performance. This corroborates previous findings demonstrating an association between greater hours of playing video games and improved spatial cognition (Feng et al., 2007; Green & Bavelier, 2003; Lin et al., 2020; Mclaren-Gradinaru et al., 2020; Murias et al., 2016). However, the strength of the association between weekly hours of video gaming and navigation performance did not significantly differ by gender, contradicting our hypothesis and suggesting that video gaming may influence navigation performance to a similar degree in men and women. This may seem at oddity with studies showing that men tend to play action-, role-play-and simulation-based video games which are more likely to have a navigational component (Quaiser-Pohl et al., 2009; Dindar. 2018, Leonhardt & Overa., 2021), and that men spend significantly longer time gaming than women (Dindar, 2018; Entertainment Software Association, 2010, 2022; Leonhardt & Overa, 2021; Lucas & Sherry, 2004; McLaren-Gradinaru et al., 2023; Ogletree & Drake, 2007; Quaiser-Pohl et al., 2006). However, we also found that those who most commonly played role-play-based video games had significantly better navigation performance than those not, suggesting that video game genre was unlikely to account for our findings (see Supplementary Table S5). Additionally, not all studies have found that men spend significantly longer gaming than women, where a recent study found that the gender difference in the spent playing video games in Scandinavian adolescents decreased from age 11 to 14, being eliminated at age 14 (Leonhardt & Overa, 2021). Therefore, it may be that specific stages of development are more prone to gender differences in the effects of video gaming on navigation ability, and these effects may well differ between countries and cultures, as has been shown for self-reported navigation skill (Walkowiak et al., 2023). Attitudes towards-and reasons for playing video games have also been found to differ by gender, where men are more likely to play male characters (Ogletree & Drake, 2007), play video games to compete (Lucas & Sherry, 2004) and play as part of a team (Dindar et al., 2018). Given that we did not examine these variables here, further investigation will be needed to determine whether differences in these variables may account for gender differences in the effect of video gaming on navigation ability.

We did find that gender was no longer significantly associated with wayfinding performance when accounting for video gaming experience. This finding aligns with a previous study where men lost their advantage in mental rotation ability whilst women gained a small advantage in spatial perspective taking ability when accounting for previous video game experience (McLaren-Gradinaru et al., 2023). Thus, the mitigation of gender differences in navigation ability in favour of males when accounting for video gaming experience may be apparent across different facets of spatial cognition, extending from mental rotation and perspective taking in this latter study to wayfinding in our study.

We considered whether our observation of the beneficial effects of video games playing was because our navigation test was in itself a video game. Future research will be useful to confirm whether real-world navigation will be enhanced by video games playing. Nonetheless, to explore this topic in the current data we examined the tutorial level (Level 1 of SHQ, which requires minimal navigation), where we reasoned participants who play a lot of video games should have an advantage taking shorter direct paths to the visible goal. However, we found no evidence for this. Rather it appears the advantage associated with video games playing is linked to the cognitively demanding navigation levels that require orientation and spatial memory. This mirrors prior research with SHQ, where we found that the tutorial level (level 1 of SHQ) was not predictive of real-world navigation in London or Paris, but rather SHQ levels that require memorising the map and navigating to the hidden multiple goals (Coutrot et al., 2019).

Finally, although we did not make a specific hypothesis about the association between phone use and navigation performance, we found a significant effect of weekly hours of phone use even when accounting for weekly hours of video gaming, age and gender, such that those spending more weekly hours using their phones were significantly worse at navigating, as indicated by their longer wayfinding distance. This finding is consistent with a range of studies that found greater hours of phone use significantly associated with poorer performance on various cognitive tasks (Chen et al., 2016; Chun et al., 2017, 2018; Hadar et al., 2017; Hovarth et al., 2020; Hutton et al., 2019; Lee et al., 2019; Paik et al., 2019; Paulus et al., 2019; Schmitgen et al., 2020; Tymofiyeva et al., 2020; Wacks & Weinstein, 2021; Wegmann et al., 2020; Weinstein et al., 2017). Here, we now show this also occurs for spatial navigation performance as tested by our measure of virtual wayfinding. Notably, the effect size for this significant association was extremely small (*f2* = 0.002). Thus if this is a true effect, it is not a major factor in predicting navigation skill.

### Limitations and future directions

While our study allowed assessment of over 800 participants on an ecologically valid (Coutrot et al., 2019) and reliable cognitive test of navigation (Couglan et al. 2020), there are a number limitations to consider that future studies could address. It would be useful to explore direct metrics for video game use and reliance on GPS from device data, rather than simply self-estimates. Self-estimates can be biassed, with studies showing different degrees of correlations with self-rating scores and performance (e.g. Garg et al., 2023). Age and the impact of different cultures may well modulate the potential impact that reliance on GPS may have on navigation, thus testing participants in a broader range of countries and age ranges would be beneficial. Future studies would also benefit from using a broader range of virtual navigation and spatial memory tests, that extending to real-world environments (Coutrot et al., 2019; He et al., 2021; Hill et al., 2023; Howard et al., 2014; Javadi et al., 2019; Patai et al., 2019). Finally, such future studies would benefit from incorporating analysis of path choices that estimate the extent to which participants are using certain strategies or heuristics for navigation (de Cothi et al., 2022; Lancia et al., 2023, McElhinney et al., 2023).

## Conclusion

Overall, our findings suggest that engaging in certain activities, such as playing video games, may indeed be beneficial to one’s navigation ability. Accordingly, this research will serve as a platform for future research looking into how actively practicing cognitive abilities during daily activities, such as playing video games, may mitigate the effect of gender and reliance on GPS on navigation performance and provide a greater understanding as to why individual differences in navigation ability may exist.

## Acknowledgements

We would like to thank all the participants who volunteered to take part in this research. This research is part of the Sea Hero Quest initiative funded and supported by Deutsche Telekom. Alzheimer’s Research UK (ARUK-DT2016-1) funded the analysis; Glitchers designed and produced the game; and Saatchi and Saatchi London managed its creation.

## Data availability

A dataset containing the preprocessed trajectory lengths and demographic information is available at https://osf.io/7nqw6/?view_only=6af022f2a7064d4d8a7e586913a1f157. Due to its considerable size (around 1 terabyte), a dataset with the full trajectories is available on a dedicated server: https://shqdata.z6.web.core.windows.net/. We also set up a portal where researchers can invite a targeted group of participants to play SHQ and generate data about their spatial navigation capabilities. Those invited to play the game will be sent a unique participant key, generated by the SHQ system according to the criteria and requirements of a specific project decided by the experimenter. https://seaheroquest.alzheimersresearchuk.org/. Access to the portal will be granted for non-commercial purposes. Future publications based on this dataset should add ‘Sea Hero Quest Project’ as a co-author.

## Code availability

The code used to produce this data is accessible at: https://osf.io/fqxzv/?view_only=ffc994fe850e4bb981fd39df0e843d80

