## Supplementary Materials for "Video gaming, but not reliance on GPS, is associated with spatial navigation performance"

| **Variable** | ***VIF*** |
|---|---|
| Age | 1.20 |
| Gender | 1.38 |
| Average GPS reliance score | 2.33 |
| Weekly hours of phone use | 1.05 |
| Weekly hours of video gaming on all devices | 2.37 |
| Highest education level achieved | 1.10 |
| Gender*Average GPS reliance score | 2.36 |
| Gender*Weekly hours of video gaming on all devices | 2.18 |
| Weekly hours of video gaming on all devices*Average GPS reliance score | 1.22 |

**Table S1. Variance inflation factor (*VIF*) values for each of the predictor variables included in the main model.**

| **Variable** | ***VIF*** |
|---|---|
| Age | 1.19 |
| Gender | 1.14 |
| Average GPS reliance score | 2.12 |
| Weekly hours of phone use | 1.03 |

| | |
|---|---|
| Highest education level achieved | 1.09 |
| Gender*Average GPS reliance score | 2.01 |

**Table S2. Variance inflation factor (*VIF*) values for each of the predictor variables included in the main model without video game experience as a predictor variable.**

| Model |
|---|
| Weighted wayfinding distance ~ Hours per week of video game play on all devices + GPS reliance score + Gender* Hours per week of video game play on all devices + Gender*GPS reliance score + Hours per week of video game play on all devices*GPS reliance score + Gender + Age + Hours per week of video game play on all devices + Hours of phone use per week + Highest education level achieved + Video game genre most commonly played + Video game platform most commonly used |

**Table S3. Model specification for the model predicting weighted wayfinding distance based on reliance on GPS and weekly video gaming hours when including the most frequently played video game genre and the most frequently used video game platform as covariates.**

| **Variable** | ***GVIF*** | ***Df*** | ***GVIF^(1/(2*Df))*** |
|---|---|---|---|
| Age | 1.253 | 1 | 1.120 |
| Gender | 1.585 | 1 | 1.259 |
| Average GPS reliance score | 2.601 | 1 | 1.613 |
| Weekly hours of phone use | 1.162 | 1 | 1.078 |

| | | | |
|---|---|---|---|
| Weekly hours of video gaming on all devices | 3.210 | 1 | 1.792 |
| Highest education level achieved | 1.118 | 1 | 1.058 |
| Most commonly played video game genre | 1.917 | 7 | 1.048 |
| Most commonly used video game platform | 2.127 | 4 | 1.099 |
| Gender*Average GPS reliance score | 2.650 | 1 | 1.628 |
| Gender*Weekly hours of video gaming on all devices | 2.612 | 1 | 1.616 |
| Average GPS reliance score*Weekly hours of video gaming on all devices | 1.176 | 1 | 1.084 |

**Table S4.** ***GVIF^(1/(2*Df))*** **values for each of the predictors included in the model predicting weighted wayfinding distance based on reliance on GPS and weekly video gaming hours when including the most frequently played video game genre and the most frequently used video game platform as covariates.**

**Model including the most commonly played video game genre and the most often used video game platform as covariates in those who game weekly**

When conducting the same multivariate model above with the most commonly used video game genre and most commonly played video game platform as covariates, average GPS reliance score remained a non-significant predictor of weighted wayfinding distance ($\beta = 0.04$, $f2 = <0.001$, $p = 0.168$, $CI = [-0.02, 0.11]$) and weekly hours of video gaming remained a significant predictor of weighted wayfinding distance $\beta = -0.07$, $f2 = 0.04$, $p = 0.040$, $CI = [-0.14, <0.001]$). Those who engaged in role-playing-based video games had significantly better navigation performance than those who did not, as indicated by their shorter wayfinding distance (*β* = -0.23, *f2* = 0.03, *p* = 0.004, *CI* = [-0.38, -0.07])(Supplementary Table S5).

| **Variable** | ***β*** | ***95% CI*** | ***t*** | ***p*** | **sig** | ***f2*** |
|---|---|---|---|---|---|---|
| (Intercept) | <0.001 | [-0.17, 0.17] | -0.01 | 0.992 | | |
| Age | 0.07 | [0.03, 0.12] | 3.42 | **0.001** | *** | 0.03 |
| Male gender | -0.08 | [-0.18, 0.01] | -1.67 | 0.095 | . | 0.03 |
| Average GPS reliance score | 0.04 | [-0.02, 0.11] | 1.38 | 0.168 | | <0.001 |
| Weekly hours of phone use | 0.05 | [0.01, 0.09] | 2.40 | **0.017** | * | 0.01 |
| Weekly hours of video gaming on all devices | -0.07 | [-0.14, <0.001] | -2.06 | **0.040** | * | 0.04 |
| Highest education level achieved | 0.07 | [-0.01, 0.16] | 1.69 | 0.091 | . | 0.01 |
| Genre - Adventure | -0.13 | [-0.29, 0.03] | -1.57 | 0.116 | | 0.03 |
| Genre - First-person shooter | -0.13 | [-0.30, 0.03] | -1.62 | 0.105 | | |
| Genre - Other | -0.10 | [-0.32, 0.12] | -0.88 | 0.382 | | |
| Genre - Role-playing | -0.23 | [-0.38, -0.07] | -2.92 | **0.004** | ** | |
| Genre - Simulation | -0.14 | [-0.32, 0.04] | -1.49 | 0.136 | | |
| Genre - Sports | 0.10 | [-0.12, 0.32] | 0.92 | 0.360 | | |
| Genre - Strategy | -0.11 | [-0.28, 0.05] | -1.34 | 0.182 | | |
| Game platform - Desktop | -0.07 | [-0.19, 0.04] | -1.25 | 0.212 | | 0.01 |
| Game platform - Laptop | -0.08 | [-0.22, 0.06] | -1.10 | 0.271 | | |

| | | | | | | |
|---|---|---|---|---|---|---|
| Game platform - Phone | 0.10 | [-0.03, 0.22] | 1.56 | 0.120 | | |
| Game platform - Other handheld device | 0.04 | [-0.13, 0.22] | 0.48 | 0.629 | | |
| Male gender*Average GPS reliances score | -0.08 | [-0.16, 0.01] | -1.84 | 0.066 | . | <0.001 |
| Male gender*Weekly hours of video gaming on all devices | <0.001 | [-0.08, 0.09] | 0.10 | 0.920 | | <0.001 |
| Average GPS reliances score*Weekly hours of video gaming on all devices | 0.05 | [0.01, 0.09] | 2.40 | **0.017** | * | 0.01 |

**Table S5. Model output when predicting weighted wayfinding distance based on reliance on GPS and weekly video gaming hours when including the most frequently played video game genre, and the most frequently used video game platform as covariates. P-values for the significant associations are highlighted in bold.** [a] **The Cohen's f2 effect sizes for the most commonly played video game genre and the most commonly used video game platform were calculated for the main effects of each of these variables as a whole rather than for each category/level of these variables.**

| **Model** |
|---|
| Weighted wayfinding distance ~ Hours per week of video game play on all devices + GPS reliance score + Gender*Hours per week of video game play on all devices + Gender*GPS reliance score + Hours per week of video game play on all devices*GPS reliance score + Gender + Age + Hours per week of video game play on all devices + Hours of phone use per week |

**Table S6. Model specification for the model predicting weighted wayfinding distance based on GPS reliance and weekly video gaming hours when using the weighted wayfinding distance from level 1 only, using video game genre as a covariate.**

| Variable | *GVIF* | *Df* | *GVIF^(1/(2*Df))* |
|---|---|---|---|
| Age | 1.24 | 1 | 1.11 |
| Gender | 1.57 | 1 | 1.25 |
| Average GPS reliance score | 2.54 | 1 | 1.59 |
| Weekly hours of phone use | 1.15 | 1 | 1.07 |
| Weekly hours of video gaming on all devices | 3.33 | 1 | 1.82 |
| Highest education level achieved | 1.12 | 1 | 1.06 |
| Most commonly played video game genre | 1.94 | 7 | 1.05 |
| Most commonly used video game platform | 2.18 | 4 | 1.10 |
| Gender*Average GPS reliance score | 2.60 | 1 | 1.61 |
| Gender*Weekly hours of video gaming on all devices | 2.70 | 1 | 1.64 |
| Average GPS reliance score*Weekly hours of video gaming on all devices | 1.16 | 1 | 1.08 |

**Table S7.** ***GVIF^(1/(2*Df))*** **values for each of the predictor variables included in the main model when using the weighted wayfinding distance from level 1 only, using video game genre as a covariate.**

| **Variable** | ***β*** | ***95% CI*** | ***t*** | ***p*** | **sig** | ***f2*** |
|---|---|---|---|---|---|---|
| (Intercept) | -0.29 | [-0.52, -0.06] | -2.43 | | | |
| Age | 0.01 | [-0.05, 0.07] | 0.39 | 0.697 | | <0.001 |
| Male gender | -0.03 | [-0.16, 0.10] | -0.46 | 0.644 | | <0.001 |
| Average GPS reliance score | 0.03 | [-0.05, 0.11] | 0.64 | 0.520 | | <0.001 |
| Weekly hours of phone use | -0.04 | [-0.09, 0.02] | -1.35 | 0.176 | | <0.001 |
| Weekly hours of video gaming on all devices | -0.03 | [-0.12, 0.06] | -0.60 | 0.550 | | <0.001 |
| Highest level of education achieved | 0.03 | [-0.09, 0.14] | 0.50 | 0.619 | | <0.001 |
| Genre - Adventure | -0.03 | [-0.25, 0.19] | -0.29 | 0.773 | | 0.01 |
| Genre - First-person shooter | 0.02 | [-0.20, 0.24] | 0.15 | 0.878 | | |
| Genre - Other | -0.05 | [-0.34, 0.25] | -0.32 | 0.749 | | |
| Genre - Role-playing | 0.10 | [-0.11, 0.31] | 0.95 | 0.340 | | |
| Genre - Simulation | 0.03 | [-0.21, 0.27] | 0.23 | 0.820 | | |
| Genre - Sports | -0.06 | [-0.34, 0.23] | -0.38 | 0.706 | | |
| Genre - Strategy | -0.04 | [-0.26, 0.18] | -0.34 | 0.735 | | |
| Game platform - Desktop | 0.23 | [0.08, 0.38] | 2.99 | **0.003** | ** | 0.02 |
| Game platform - Laptop | 0.13 | [-0.05, 0.32] | 1.39 | 0.165 | | |

| | | | | | | |
|---|---|---|---|---|---|---|
| Game platform - Phone | 0.17 | [0.01, 0.34] | 2.07 | **0.039** | * | |
| Game platform - Other handheld device | 0.21 | [-0.02, 0.44] | 1.81 | 0.072 | . | |
| Male gender*Average GPS reliances score | -0.06 | [-0.17, 0.05] | -1.10 | 0.272 | | <0.001 |
| Male gender*Weekly hours of video gaming on all devices | -0.05 | [-0.16, 0.07] | -0.80 | 0.425 | | <0.001 |
| Average GPS reliances score*Weekly hours of video gaming on all devices | -0.05 | [-0.10, 0.01] | -1.74 | 0.082 | . | <0.001 |

**Table S8. Model output when predicting weighted wayfinding distance based on reliance on GPS and weekly video gaming hours when using the weighted wayfinding distance from level 1 only, using video game genre as a covariate. P-values for the significant associations are highlighted in bold.** [a] **The Cohen's f2 effect sizes for the most commonly played video game genre and the most commonly used video game platform were calculated for the main effects of each of these variables as a whole rather than for each category/level of these variables.**

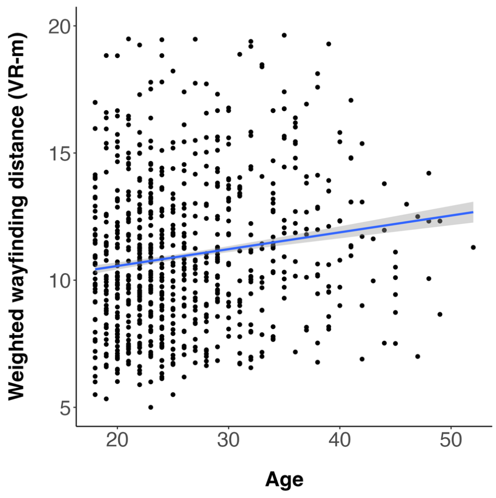

A)

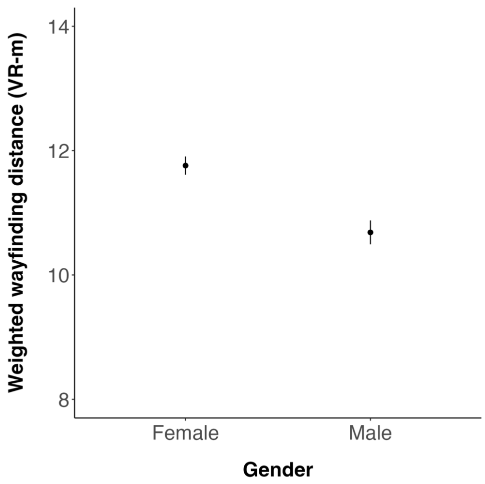
Weighted wayfinding distance (VR-m)
14
12
10
8
Female
Male
Gender

B)

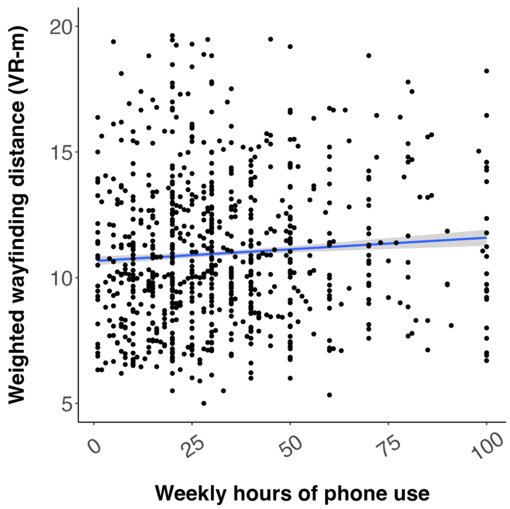
Weighted wayfinding distance (VR-m)
20
15
10
5
0
25
50
75
100
Weekly hours of phone use

C)

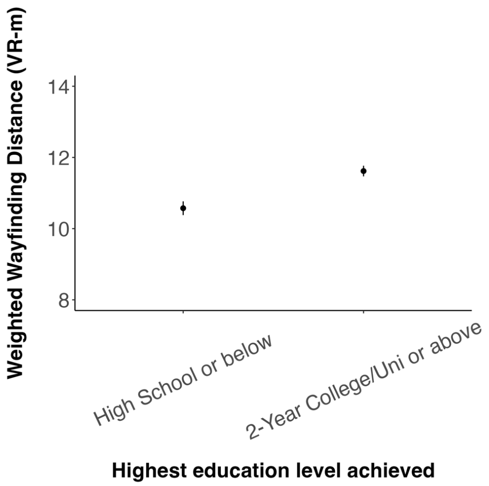


D)

**Figure S1. (A-F) Associations between each of the demographic variables and weighted wayfinding distance.**

**(A,C) Blue line indicates the mean wayfinding distance across game levels across participants. Grey shading surrounding the blue line indicates the standard error of the mean corresponding to this wayfinding distance. Data points indicate the mean wayfinding distance across game levels for an individual participant. VR-m = virtual reality metres.**

**(B,D) Data points represent the mean wayfinding distance across game levels across participants. Bars represent the standard error of the mean corresponding to this wayfinding distance. VR-m = virtual reality metres.**

<u>Assessing the internal reliability of the NSQ, GPS reliance scale and the SHQ weighted wayfinding distance</u>

*NSQ*

Exploratory factor analysis revealed that the NSQ could be explained by 3 factors, as determined by a scree plot which showed that 3 factors had eigenvalues greater than or equal to 1 (Zwick & Velicer, 1986)(Figure S1). When we looked at the loadings of each item of the NSQ onto each of these 3 factors, including only items with a loading greater than 0.4, the first factor only consisted of items relating to navigation ability (questions 2, 3, 4, 12, 13 and 14) with the exception of question 13, whilst the second factor consisted of items only relating to navigation strategy (questions 5, 6 and 11). The 3rd factor did not have any items with a loading >0.4, which supported the notion that the NSQ could be split into a strategy and ability subscale (Table S2A). However, the cumulative variance explained by factors 1 and 2 was 25%, suggesting that these factors only weakly accounted for the NSQ scores (Table S2B). McDonald's omega factor saturation scores for the total, ability and strategy subscales of the NSQ were 0.74, 0.82 and 0.56 respectively (Table S2C). Split-half permutation reliability analysis indicated the total and ability subscales of the NSQ had a Spearman-Brown correlation value of -0.02 and 0.17, whilst the strategy subscale had too few items to calculate a reliability score (Table S2D). A chi-squared test indicated that the 3 questions of the NSQ strategy subscale were significantly dependent on one another, indicating that there was a relationship between the questions in the NSQ strategy subscale (question 5 vs question 11: $\chi 2 = 30.583$, $p < 0.001$, $V = 0.136$, question 5 vs question 6: $\chi 2 = 357.660$, $p < 0.001$, $V = 0.466$, question 6 vs question 11: $\chi 2 = 284.941$, $p < 0.001$, $V = 0.416$)(Table S2E). Cronbach's alpha scores for the total, ability and strategy subscales were 0.37, 0.23 and 0.50 respectively (Table S2F). These findings would suggest that the NSQ is only a weak measure of the extent to which somebody relies on map-based strategies in the real world. We thus decided not to include the NSQ in our analysis when looking at how the self-report questionnaire responses are associated with wayfinding distance.

*GPS reliance scale*

Split-half permutation reliability analysis indicated the GPS reliance scale had a Spearman-Brown correlation value of 0.39 (Table S2D). McDonald's omega factor saturation score for the GPS reliance scale was 0.79 (Table S2E). The Cronbach's alpha score for the GPS reliance scale was 0.80 (Table S2F).

*Weighted wayfinding distance*

Split-half permutation reliability analysis indicated the GPS reliance scale had a Spearman-Brown correlation value of -0.37 (Table S2D). McDonald's omega factor saturation score for the GPS reliance scale was 0.27 (Table S2E). The Cronbach's alpha score for the weighted wayfinding distance across the game levels (11, 32, 42 and 68) was 0.64 (Table S2F).

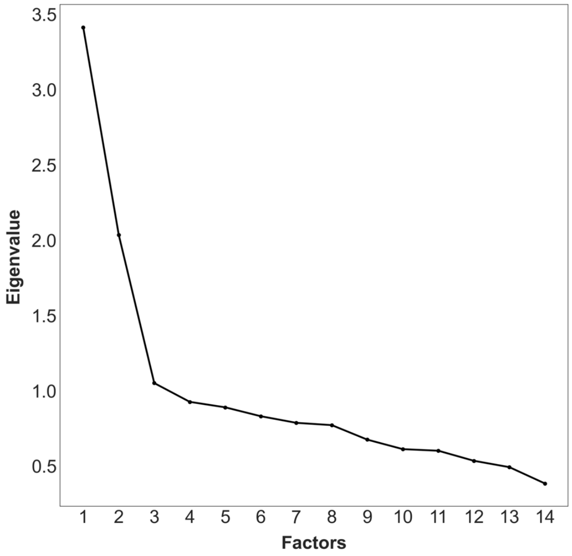


**Figure S1. Scree plot showing the number of factors against the corresponding eigenvalue.**

| Factor 1 | | Factor 2 | |
|---|---|---|---|
| **Question** | **Loading** | **Question** | **Loading** |
| response2q | 0.71 | response11q | 0.60 |
| response3q | 0.68 | response6q | 0.51 |
| response4q | 0.68 | response5q | 0.47 |
| response13q | 0.62 | | |
| response12q | 0.54 | | |
| response14q | 0.44 | | |

A)

| | Factor 1 | Factor 2 | Factor 3 |
|---|---|---|---|
| SS Loadings | 2.54 | 1.00 | 0.96 |
| Proportion Var | 0.18 | 0.07 | 0.07 |
| Cumulative Var | 0.18 | 0.25 | 0.32 |

B)

| Scale | McDonald's total omega (*g*) |
|---|---|

| | |
|---|---|
| Complete NSQ questionnaire | 0.74 |
| NSQ ability subscale | 0.81 |
| NSQ strategy subscale | 0.55 |
| GPS reliance scale | 0.79 |
| Weighted wayfinding distance | 0.27 |

C)

| **Scale** | **Spearman-Brown coefficient (*r*)** |
|---|---|
| Complete NSQ questionnaire | 0.10 |
| NSQ ability subscale | -0.07 |
| NSQ strategy subscale | NA (too few subscale items) |
| GPS reliance scale | 0.39 |
| Weighted wayfinding distance | -0.37 |

D)

| **Comparison** | **$\chi 2$** | **Df** | ***p*** | **Cramer's *V*** |
|---|---|---|---|---|
| q5 vs q11 | 30.6 | 2 | **<0.001** | 0.14 |

| | | | | |
|---|---|---|---|---|
| q5 vs q6 | 357.7 | 2 | **<0.001** | 0.47 |
| q6 vs q11 | 284.9 | 2 | **<0.001** | 0.42 |

E)

| | **Alpha** | ***95% CI*** |
|---|---|---|
| GPS reliance scale | 0.80 | [0.78, 0.82] |
| NSQ ability subscale | 0.22 | [0.13, 0.30] |
| NSQ strategy subscale | 0.49 | [0.43, 0.55] |
| NSQ total questionnaire | 0.35 | [0.29, 0.42] |
| Weighted wayfinding distance | 0.62 | [0.07, 0.26] |

F)

**Table S9 (A-F). Exploratory factor analysis and internal reliability assessment outcomes for the NSQ, GPS reliance scale and SHQ weighted wayfinding distance scores using various psychometric measures.**

(A) Loadings of each item (question) onto each of the 3 factors retained in the exploratory factor analysis. Items with a loading > 0.4 were retained.

(B) Cumulative and proportional variance explained by the 3 factors retained in the exploratory factor analysis.

(C) McDonald's omega factor saturation scores for the NSQ full questionnaire, NSQ ability subscale, NSQ strategy subscale, GPS reliance scale and weighted wayfinding distance across the 5 levels (1, 11, 32, 42, 68).

(D) Split-half permutation reliability testing for the NSQ full questionnaire, NSQ ability subscale, NSQ strategy subscale, GPS reliance scale and weighted wayfinding distance across the 5 levels (1, 11, 32, 42, 68).

(E) Chi-squared analysis comparing each item (questions 5,6 and 11) in the NSQ strategy subscale.

(F) Cronbach's alpha scores for the NSQ full questionnaire, NSQ ability subscale, NSQ strategy subscale, GPS reliance scale and weighted wayfinding distance across the 4 game levels (11, 32, 42, 68).
